# Risk assessment for condylar stress fracture in elite racing Thoroughbreds using standing computed tomography-based virtual mechanical testing

**DOI:** 10.1101/2025.05.13.653480

**Authors:** Nicola L. Brown, Soroush Irandoust, Elleana J. Thom, R. Christopher Whitton, Corinne R. Henak, Peter Muir

## Abstract

**Background:** Condylar stress fracture of the third metacarpal bone is a common catastrophic musculoskeletal injury in Thoroughbred racehorses. Condylar stress fracture is associated with parasagittal groove (PSG) subchondral osteolysis. Standing computed tomography (sCT) imaging allows for sensitive identification of this fatigue-induced structural change, an early form of subchondral bone injury (SBI), but there is not yet an objective method for identifying racehorses at heightened risk of condylar stress fracture.

**Objectives:** To estimate PSG first principal strain in elite Thoroughbred racehorses that have undergone subjective risk assessment using sCT fetlock screening.

**Study design:** Retrospective clinical study.

**Methods:** We used fetlock sCT images from 9 thoracic limbs from 7 Thoroughbred racehorses. A tuned, validated 3D finite element (FE) analysis approach was used as a virtual mechanical test to estimate PSG first principal strain in the distal third metacarpal bone (MC3) from these joints. Virtual mechanical testing results were compared with a subjective clinical risk assessment using an established screening approach by Racing Victoria.

**Results:** MC3 condyles with PSG SBI consistently and significantly displayed increased levels of first principal strain throughout the PSG. We found focal strain concentrations associated with the anatomical location of SBI compared to condyles with no evidence of SBI in the PSG. Clinical diagnosis of SBI with focal osteolysis, FE-predicted strain elevation, and clinical risk assessment were concordant with recall and precision of 0.86.

**Main limitations:** The sample size was small, and our virtual mechanical testing protocol does not account for whole-joint physiology.

**Conclusions:** Risk assessment through screening with sCT is an established approach to injury prevention in racing Thoroughbreds. Concordance of a current clinical risk assessment approach by Racing Victoria with objective FE analysis of principal strain in sites of SBI in the present study suggests 3D FE analysis using a validated pipeline is an important new approach for routine assessment of risk of MC3 condylar stress fracture in Thoroughbred racehorses.

## 1 INTRODUCTION

Catastrophic musculoskeletal injuries (CMI) are a leading cause of euthanasia in Thoroughbred racehorses, with a pooled incidence of 1.17 CMI per 1,000 race starts worldwide from 1990 to 2017.^1^ Condylar stress fracture of the third metacarpal/tarsal bone (MC3/MT3) is a prevalent example of CMI and frequently results in euthanasia.^1,2^ Catastrophic injuries also increase the risk of jockey injury.^3^

Condylar stress fractures of the MC3 occur due to an accumulation of microdamage^4^ that compromises the mechanical integrity of the distal end of the bone, most commonly on the palmar surface,^5,6^ due to the repetitive high loads the bone experiences during galloping.^7^ Subchondral trabecular bone becomes sclerotic in response to the repetitive loading, and under low strain conditions the damaged bone undergoes remodeling to repair the microcracks and restore mechanical integrity, resorbing damaged bone and replacing it with new bone.^8^ When the bone is excessively subjected to high loads, such as the cyclic loads associated with galloping, microdamage accumulates faster than it can be fully repaired through remodeling, resulting in damage accumulation, intense targeted remodeling, and focal resorption. Such changes are commonly observed in the PSG subchondral bone^8,9^ This type of subchondral bone injury (SBI) is visible as focal subchondral bone resorption in the PSG with standing CT (sCT) imaging.^10^

sCT is a highly sensitive method of detecting fatigue-induced structural changes in subchondral bone. Such changes in the PSG are associated with condylar fracture and local elevated strain when the articular surface is loaded.^5,11^ However, interpretation of the significance of these lesions is subjective and debated among clinicians.^10^ Therefore, an objective method of assessing risk of incipient fracture is needed. Virtual mechanical testing of *ex vivo* MC3 bones by 3D sCT-based finite element (FE) analysis predicted elevated PSG strain in bones with PSG SBI.^12^ Incorporation of the different bone material properties associated with sub volumes of sclerotic and lytic bone increased the accuracy of PSG strain prediction. This approach has been validated using *ex vivo* bone specimens but has not yet been applied to live Thoroughbred racehorses.

We hypothesized that 3D FE analysis of *in vivo* limbs of actively racing and training elite Thoroughbred racehorses would predict elevated strain in the PSG with presence of SBI, thereby indicating elevated risk of condylar stress fracture. In addition, risk assessment scores made by clinicians were compared with FE-predicted PSG strain to investigate alignment of the current subjective sCT-based clinical risk assessment with the objective assessment using virtual mechanical testing.

## 2 MATERIALS AND METHODS

### 2.1 Diagnostic imaging and sample population

Four fetlock sCT scans were performed at the University of Melbourne Equine Centre using an Equina^®^ sCT scanner with an exposure of 160 kVp and 8 mA, with 1 mm slice thickness. An electron density phantom (model 062M, CIRS Inc, Arlington, VA) with 4 plugs of 200, 800, 1250, and 1500 *mg*_*HA*_/*cm*^3^was scanned asynchronously for calibration of CT density (*ρ*_*CT*_, **Supplementary Figures S1-2**). Nine thoracic limbs from seven elite Thoroughbred horses training and racing in the 2021, 2022, and 2023 Spring Racing Carnival events under the oversight of Racing Victoria were selected to include thoracic limbs with and without PSG SBI (**Table 1**). The horses ranged in age from 3yo to 7yo. There were six geldings and one mare. Digital radiographs were also available for qualitative comparison with sCT for detection of structural changes in the PSG in two horses. Flexed dorsopalmar digital radiographs were made 74 and 43 days before sCT was performed for Horses #2 and #4, respectively.

**Table 1.** Risk assessment for Thoroughbreds racing in Melbourne, Australia.

| ID | Clinical risk scores | Limb | LC PSG | MC PSG |
| --- | --- | --- | --- | --- |
| 1 | 1 (0, 1, 1) | LF | CTRL | CTRL |
| 2* | 2 (1, 2, 2, 2) | LF | CASE | CASE |
|  |  | RF | CASE | CTRL |
| 3 | 1 (0, 1, 1) | RF | CTRL | CTRL |
| 4 | 0 (0, 0, 0) | LF | CTRL | CTRL |
| 5 | 2 (1, 2, 2) | LF | CTRL | CASE |
|  |  | RF | CASE | CTRL |
| 6 | 2 (1, 2, 2) | RF | CASE | CTRL |
| 7 | 2 (2, 2, 2) | LF | CASE | CTRL |

**Note.** CASE and CTRL indicate existence and absence of parasagittal groove subchondral bone injury, respectively. *Racing Victoria Risk Assessment scale* was 0 – Standard level of injury risk, 1 – Moderately elevated level of injury risk, 2 – Heightened level of injury risk. Risk assessment was based in review of standing computed tomography diagnostic imaging for structural change by a panel of three expert reviewers. LF – left thoracic limb, RF – right thoracic limb, LC – lateral condyle, MC – medial condyle, PSG – parasagittal groove. \*This horse was reviewed by four experts.

### 2.2 Clinical risk assessment

Assessment of structural changes in the fetlock was undertaking from sCT images by a panel of 3 or 4 expert veterinarian consultants. For Horse #2 with PSG SBI and #4 without PSG SBI, digital radiographs were also available (**Table 1**). Expert veterinarian consultants provided assessment of risk of injury for each horse as standard risk (0), intermediate risk (1), and heightened risk (2) (**Table 1**). The associated heightened injury risk was only due to PSG lesions, as no other concerning fatigue injury was present in these horses.

### 2.3 Virtual mechanical testing

Our virtual mechanical testing pipeline has been described in detail previously^12^ and is summarized here. The distal MC3 bone was segmented using Mimics (v.26), and based on the density threshold of HU=1,200 (*ρ*_*CT*_ = 1,055.75 *mg*_*HA*_/*cm*^3^), the sclerotic subchondral trabecular bone in the palmar aspect (HU≥1,200), and the lytic regions in the PSG (HU≤1,200, if present) were isolated. This threshold was different from what was used in our previous study to capture the sclerotic subchondral bone properly. Thresholds of HU=1,250 and HU=1,150, and HU=1,100 and HU=1,200 were used for segmentation of the lytic and sclerotic volumes, respectively, to assess the sensitivity of the predicted PSG strain to these thresholds. It is important to note that sCT scans acquired at different times would be expected to have different calibration equations resulting in variation in HU values for matched bone mineral density values, however that variation is less than 45 HU,^13^ thus the results of the sensitivity study are informative for the error expected from scanning at different times.

Three dimensional distal MC3 and the sclerotic and lytic volumes were then smoothed and imported in 3-matic (v.18), and a uniform triangular surface mesh with 0.25 mm edge length was created for all surfaces. Anatomical planes were established by first creating the transverse plane by finding its normal vector as the axis of a cylinder fit to the distal MC3. The sagittal plane was created normal to the transverse plane, and through two points on the palmar and dorsal aspects of the sagittal ridge. The frontal plane was then defined perpendicular to the transverse and sagittal planes. The distal 2.5 in (6.35 cm) of the MC3, measured from the most distal part of the sagittal ridge along the long axis of the bone, was separated for virtual mechanical testing. For each condyle, a 3 mm strip in the dorsopalmar direction centered around the PSG on the palmar aspect of the condyle, covering the 90 degrees between a frontal plane through the transverse ridge and a transverse plane through the most palmar end, was marked to monitor PSG strain.

These 3 mm strips were split into six equally angled regions of 15 degrees each for regional strain analysis (**Figure 1**). Additionally, the area between the transverse ridge and the most palmar aspect in the distopalmar direction, and half of the length between the most lateral/medial aspect and the sagittal ridge in the lateromedial direction was marked for loading (**Figure 1**).

**Figure 1.**
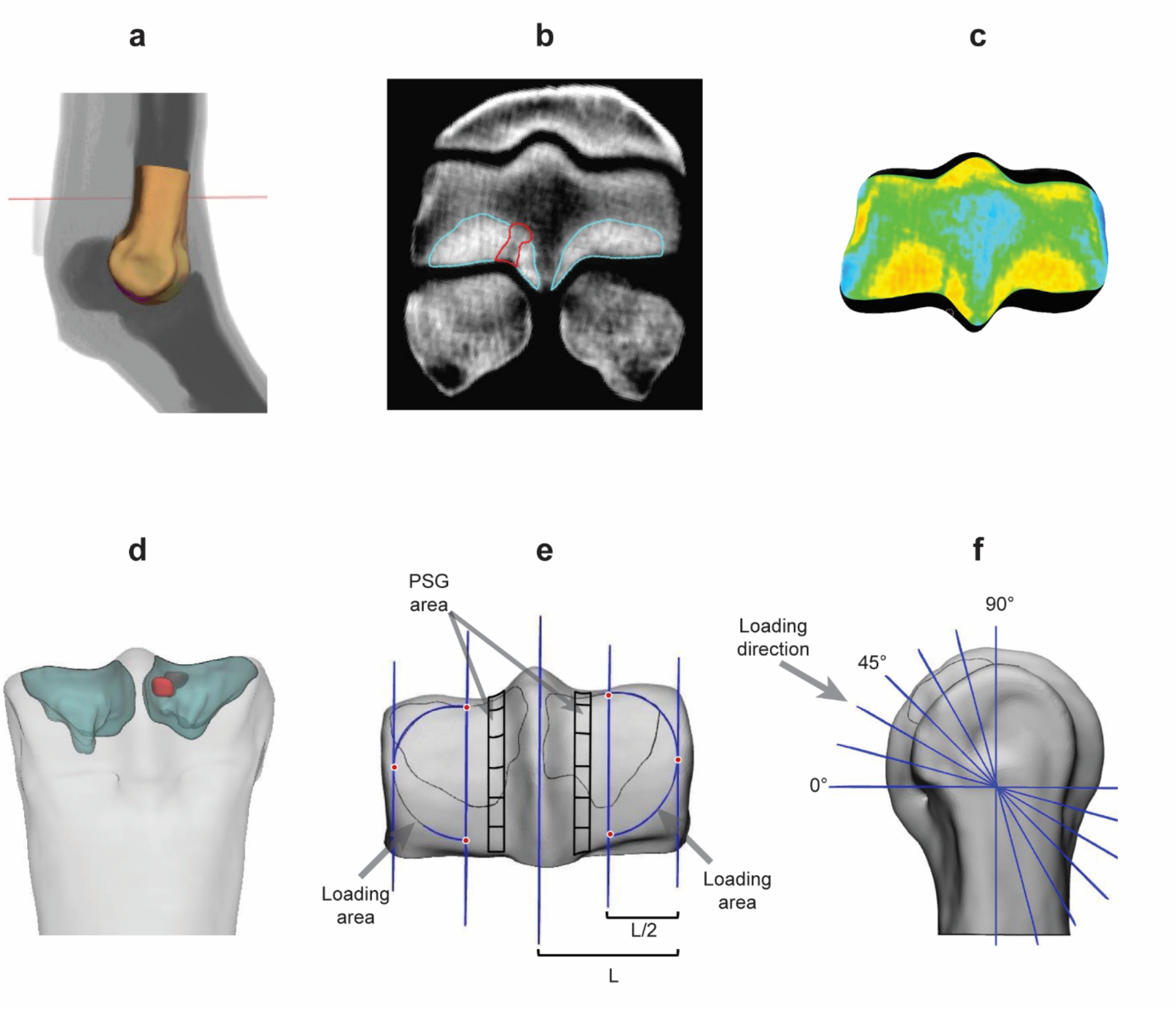
3D segmentation of the distal third metacarpal bone from the standing CT image set (a) and subsequent segmentation of the subchondral sclerotic bone (cyan) and parasagittal groove (PSG) lysis (red), if present (b). The ratio of the volume of each sclerotic region to the volume of its corresponding condyle was used to calculate the adaptive factor. An adaptive factor was applied to the regions of sclerosis. CT density was mapped in the model at each element individually (c). In the 3D model (the dorsopalmar view is shown in panel d) 3 mm wide strips centered at the PSG of both condyles were defined and divided into six equally angled regions in 15 degrees intervals from the most palmar end to the transverse ridge (e, f). The loading area was created with a straight edge in the dorso-palmar direction at the half distance between the sagittal ridge and the most lateral/medial aspect (*x*), and a curve fit on the three points shown on the image.

The 3D distal MC3s were imported in FEBio Studio (v.2.6) and discretized into 0.25 mm linear tetrahedral elements for the sclerotic and lytic regions, and 1 mm linear tetrahedral elements for the remainder of the distal MC3 bone. Elementwise (heterogenous) isotropic linear elastic material properties were assigned with Young’s modulus tuning in the sclerotic and lytic regions according to our validated methodology.^12^ This requires an adaptive factor defined for the sclerotic region (−1.00<*A*_*SCL*_<0.50), and a damage factor for the lytic region (*D*_*LYS*_=0.65). Element-wise ash density (*ρ*_*ash*_) was calculated from CT density (*ρ*_*ash*_), and Young’s modulus was calculated for the sclerotic regions, PSG SBI regions (if present), and the rest of the distal MC3 individually. A constant Poisson’s ratio of 0.3 was used.

The proximal surface of the model was constrained and a uniformly distributed 7.5 kN load with convergence required at 2.5 kN increments was applied to the previously marked area for loading in each condyle at 60 and 30 degrees with respect to the frontal and transverse planes respectively, replicating prior *ex vivo* mechanical testing loading conditions.^5^ FE-predicted strain in 6 regions of the PSG was examined for each condyle (**Figure 1**) and strain levels were compared between condyles with and without sCT-detectable PSG-SBI, cases and controls respectively.

### 2.5 Statistical analysis and risk classification

Mean PSG first principal strain and was compared between CASE and CTRL condyles using the Wilcoxon Rank-Sum test. The clinical fracture risk assessments were qualitatively compared between CASE and CTRL condyles as well. Next, a logistic regression model was built as a risk classifier (low/high risk) based on the CASE/CTRL assessments and the associated FE-predicted PSG strain, with a threshold of 0.5. The performance of the classifier was assessed using the leave-one-out cross validation (LOOCV) method. The power of the classifier was calculated with a significance threshold of <0.05 and resampling strain for 1,000 simulations. The effect of the sample size on the classifier power was estimated with resampling strain with sample sizes at n = 5 intervals ranging from n = 10 to n = 50.

## 3 RESULTS

Condyles with PSG SBI showed elevated first principal strain in the PSG and focally around the SBI lesion compared to CTRL condyles (**Figure 2**). The PSG SBI and the associated focal elevation of strain were most obvious in the transverse oblique slice in both the sCT images and the 3D FE model. Qualitative comparison of the FE-predicted strain distribution on the joint surface also showed elevated PSG strain in the CASE condyles compared with the CTRL condyles. In addition to the palmar aspect of the PSG of the CASE condyles, strain was elevated in the dorsal aspect in the grooves, evident in the transverse oblique view. With respect to diagnostic imaging, the flexed dorsopalmar digital radiographs did not reveal evidence of PSG SBI and subchondral osteolysis in the horse with SBI. The sCT image sets and FE-predicted strains plots for the rest of the limbs are shown in **Figure S3** to **Figure S9**.

**Figure 2.**
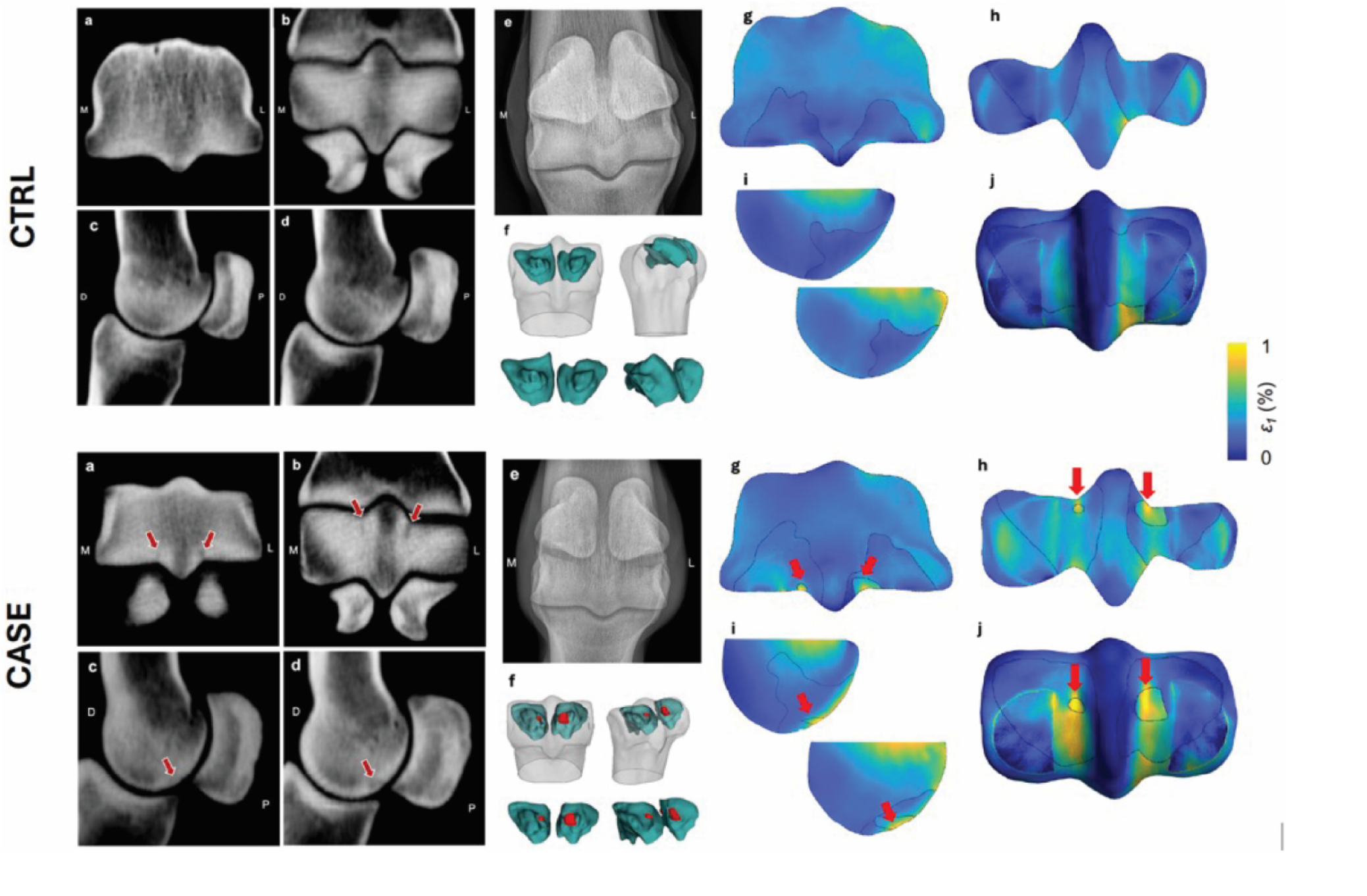
sCT images of the left metacarpophalangeal joint for a horse with control (CTRL) condyles (Horse #4 LF) and a horse with biaxial parasagittal groove (PSG) subchondral bone injury (SBI) (CASE, Horse #2 LF. Frontal oblique (a), transverse oblique (b) and two parasagittal views at the lateral and medial PSGs (c, d) from the left metacarpophalangeal joint sCT images, are shown. The flexed dorsopalmar digital radiograph (e) of each limb is shown for comparison with the sCT slices. No PSG osteolysis is evident on the radiographs. The 3D geometry of the FE model with subchondral lytic, if present, and sclerotic volumes (f). The FE-predicted first principal strain is shown in the same sCT slices (g, h, and i) and on the joint surface (j). Red arrows indicate PSG SBI and the associated elevated strain. L, M, D, and P labels on the sCT images indicate lateral, medial, dorsal, and palmar, respectively.

First principal strain was increased in the CASE condyles compared with the CTRL condyles (*p*<0.001, **Figure 3**). In the medial PSGs, first principal strain peaked close to the transverse ridge in both CASE and CTRL condyles. In the lateral PSGs, strain was lower from the transverse ridge toward the most palmar/proximal end in the CTRL condyles, while it peaked around the lesion in the CASE condyles. The threshold values used for segmentation of the sclerotic and lytic volumes had a small influence on the predicted PSG strain (**Figure S10**). All horses with CASE condyle(s) were clinically assessed to have heightened risk of injury from condylar stress fracture.

**Figure 3.**
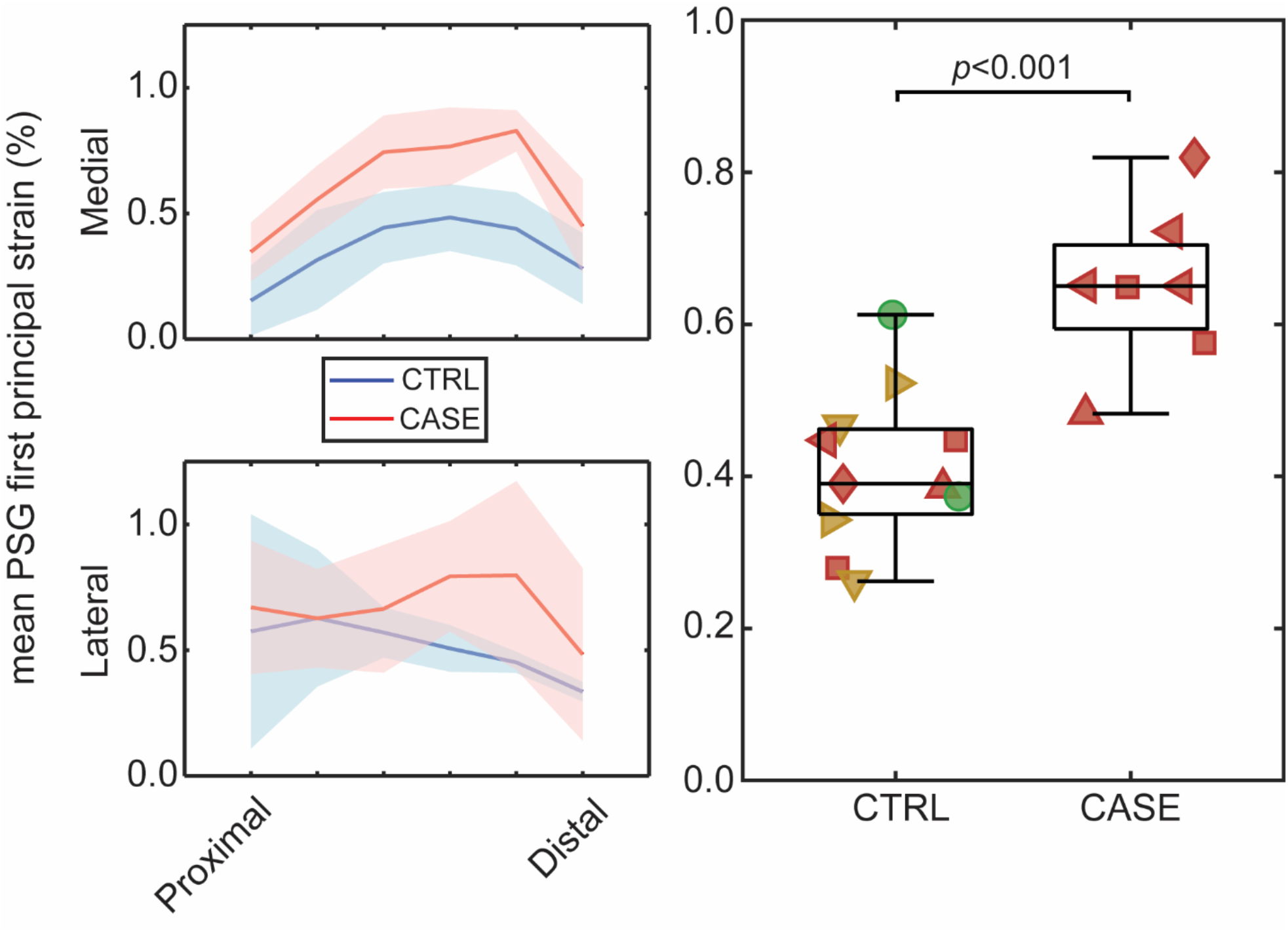
Mean PSG first principal strain was qualitatively higher in both medial and lateral sides of the CASE condyles. Strain peaked closer to the transverse ridge in the medial side in both CASE and CTRL condyles, and in the lateral side of the CASE condyles. The shaded area indicates the 95% confidence interval. In the box plot, condyles (two or four) of the same horse have the same shape, and the green, yellow, and red colors indicate standard, moderate, and heightened risk of injury, respectively, based on the median clinical scores shown in Table 1. Mean PSG strain was higher in the CASE condyles (*p*<0.001), and all horses with CASE condyles were clinically assessed to have concerning injury risk. All the red datapoints in the CTRL group had a contraaxial CASE condyle.

The logistic regression classifier had a power of 0.4 and recall and precision scores of 0.86 with one false positive and one false negative prediction. The model’s power is estimated to reach 0.80 at a sample size of n=30 (**Figure S11**).

## 4 DISCUSSION

In this report, we evaluated use of a virtual mechanical testing approach that has been validated against experimental data.^5,12^ The fetlock joints of the horses we studied with PSG SBI had lesions that are typical of affected horses.^4,14,15^ As previously described,^10^ Horse #2 with PSG SBI in the present study was readily identified by sCT, but not by digital radiography using the flexed dorsopalmar view. This aligns with past work where reliance on fetlock digital radiography likely underestimates risk of serious injury from condylar stress fracture in racing Thoroughbreds.^10^

The results of this study support our hypothesis that FE-predicted PSG strain was larger in condyles with PSG SBI compared to those without. The elevated strain is indicative of mechanical compromise associated with SBI in the PSG,^5,15^ and may be indicative of elevated risk of condylar stress fracture, as strain levels exceeding the bone’s capability is the cause of the condylar stress fracture. These observations are also supported by the reduction in catastrophic injuries that has occurred since routine sCT screening and subjective review of sCT structural changes was used for clinical risk assessment by Racing Victoria in 2021 until now.^16^

Condylar stress fracture risk has been linked to subchondral plate functional adaptation by bone modeling.^17,18^ Subchondral sclerosis acts to increase the bone volume fraction of the subchondral plate to better resist the high cyclic loads associated with racing and we found it necessary to consider this feature specifically in our virtual mechanical testing pipeline to properly estimate first principal strain in the MC3 bone end.^12^ Similarly, the dimensions of the PSG subchondral focal osteolysis lesion are thought to cause a reduction in MC3 stiffness^15^ and we found that consideration of the subchondral osteolytic volume was important to our virtual mechanical testing pipeline.^12^ Contralateral PSG subchondral lucencies and associated fatigue injury are known to be associated with catastrophically injured horses,^17^ and we found that the bones of horses with this lesion had consistently elevated first principal strain in the affected PSG.

Fetlock MC3 SBI was most easily visible on the transverse oblique slice of sCT images, supporting the use of this view as a standard screening tool as in past work.^10,19^ In the horses of this work, we identified good concordance of case control classification between the 3D FE virtual mechanical testing and the subjective clinical risk assessment through Racing Victoria’s Injury Prevention Program. The good concordance in these results is an important finding as past work suggests clinician observers are generally better at detecting and assessing structural changes from fetlock diagnostic imaging than providing a rating for risk of imminent injury.^10^

The precision and recall of the logistic regression classifier were greater than 0.80, with one false positive and one false negative prediction between the FE-predicted strain and the clinical risk scores suggesting FE analysis for virtual mechanical testing is clinically useful for identification of horses with heightened injury risk. Both false positive and false negative predictions are potentially concerning. On the one hand, allowing horses with heightened injury risk to continue to race will lead to ongoing serious or fatal injury and concerns regarding Thoroughbred and jockey welfare. Furthermore, removing horses from racing unnecessarily is financially concerning. In the horses of the present study, a fetlock of one CTRL horse had FE-predicted PSG strain that approximated the CASE group and a fetlock of one CASE horse had FE-predicted PSG strain that approximated the CTRL group. This certainly suggests more data are needed to robustly train the logistic regression classifier to further improve the precision and recall of risk assessment predictions so that high-risk CASE horses can be accurately identified for personalized care. Implementation of our virtual mechanical testing approach in a longitudinal clinical study that follows horses over time would also further affirm predictive accuracy of the case/control classifier.

In the horses of the present study, we identified qualitative differences in the pattern of first principal strain between the lateral and medial PSGs. In the lateral PSGs, strain decreased from the transverse ridge toward the most palmar/proximal end in the CTRL condyles while it peaked around the lesion in the CASE condyles, whereas in the medial PSGs strain increased distally in both CASE and CTRL condyles. The presence of some degree of PSG strain elevation in all lateral condyles, as compared to medial condyles, may suggest a mechanism by which SBI preferentially develops in the PSG of the lateral condyle. This fits with fracture epidemiology knowledge, as the incidence of lateral condylar stress fracture is higher than that of medial condylar stress fracture.^20^

There were several limitations to this study. Our analysis was based on a relatively small sample of elite Thoroughbred racehorses that may not cover all potential clinical features or range of lesion severities. In the two horses with digital radiographs, the radiographic images were not made at the same time as the sCT images. It is possible that analysis of a larger sample of Thoroughbreds may identify racehorses with discordant findings between clinical risk assessment and 3D FE analysis. However, discovery of such cases would likely inform development of a comprehensive condylar stress fracture risk assessment approach and ultimately improve the accuracy of the classifier. The risk assessment approach described in Table 1 was developed from the expertise of Racing Victoria and the consultants that make up their imaging review panel. While initial clinical outcomes are promising, further evaluation of such assessments is still needed to fully validate the approach as a clinically useful and accurate predictor. One of the specific challenges in further *in vivo* longitudinal clinical studies is that allowing horses assessed as having a high risk of imminent serious injury to continue to participate in racing and monitoring these animals to see if a stress fracture develops is not ethical and would not be viewed favorably by society. Use of virtual mechanical testing assessment of sCT images has great potential to address such a challenge. An additional challenge stems from assigning region-wide damage and adaptive factors to sclerotic and lytic regions, which do not account for the gradient of damage which is likely more severe closer to the articular surface. However, the threshold values we used for segmentation of the sclerotic and lytic volumes had a relatively small influence on predicted PSG principal strain, suggesting that damage gradients may not improve predictions. An additional limitation on the material modeling is that the post-yield behavior of the bone is not included and models, therefore, cannot predict fracture load, although this does not preclude its clinical utility as a CASE/CTRL classifier. Future studies that analyze variation in fatigue damage within sclerotic and lytic regions as well as 3D FE models that integrate physiological stress fracture are needed to better encompass accurate material properties of sclerotic and lytic regions of bone and better determine whether risk of fracture is truly imminent. The virtual mechanical testing we describe does not account for whole-joint physiologic loading, which should also be explored in future work. Incorporation of the proximal sesamoid bones and the proximal phalange as well as tendons, ligaments and muscles would create a model that more closely reflects true *in vivo* physiologic loading.

## 5 CONCLUSION

The sCT-based virtual mechanical testing approach described in this report provides an objective, non-invasive method of assessing imminent risk of condylar stress fracture in Thoroughbred racehorses and aligns well with the risk assessment grading system developed by Racing Victoria that uses subjective assessment of sCT structural change. Continued application of this approach to more bones will provide insight into interpretation of structural changes visible with sCT imaging, determining whether the shape of SBI or relative volumes of SBI and sclerosis have an impact on strain prediction and therefore fracture risk. Mandatory sCT screening and risk assessment before racing is a clinically feasible approach that promises accurate identification of horses with high imminent risk of bone fracture and enables removal of high-risk horses from racing for personalized care, ultimately reducing the incidence of condylar stress fracture, euthanasia in Thoroughbred racehorses, and injury of jockeys.^3^

## Author contributions

**Nicola L. Brown:** Methodology, Investigation, Writing -original draft, Visualization. **Soroush Irandoust:** Conceptualization, Methodology, Validation, Formal analysis, Investigation, Writing -original draft, Writing -review & editing, Visualization, Supervision. **Elleana J. Thom:** Visualization. **R. Christopher Whitton:** Conceptualization, Resources, Writing -review & editing, Funding acquisition. **Corinne R. Henak:** Conceptualization, Methodology, Formal analysis, Resources, Writing -original draft, Writing -review & editing, Supervision, Funding acquisition. **Peter Muir:** Conceptualization, Methodology, Formal analysis, Resources, Writing -original draft, Writing -review & editing, Supervision, Project administration, Funding acquisition.

## Ethical animal research

Not applicable. All data were collected from Racing Victoria’s clinical injury prevention program.

## Acknowledgements

The authors are very grateful to Racing Victoria for providing the sCT DICOM image sets and clinical risk assessment data used in this study.

