## Supplementary File for "Risk assessment for condylar stress fracture in elite racing Thoroughbreds using standing computed tomography-based virtual mechanical testing"

Manuscript for *Equine Veterinary Journal*

May 2025

Nicola L. Brown[1,#], Soroush Irandoust[1,2,#], Elleana J. Thom[1,2],

R. Christopher Whitton[3], Corinne R. Henak[2,4,5], Peter Muir[1*]

[1]Department of Surgical Sciences, University of Wisconsin-Madison, Madison, WI, USA

[2]Department of Mechanical Engineering, University of Wisconsin-Madison, Madison, WI, USA

[3]Department of Veterinary Clinical Sciences, Melbourne Veterinary School, Faculty of Science, University of Melbourne, Werribee, Victoria, Australia

[4]Department of Biomedical Engineering, University of Wisconsin-Madison, Madison, WI, USA

[5]Department of Orthopedics & Rehabilitation, University of Wisconsin-Madison, Madison, WI, USA

[#]Contributed equally to this study

**Keywords**: standing computed tomography, condylar stress fracture, Thoroughbred racehorses, finite element analysis, third metacarpal bone.

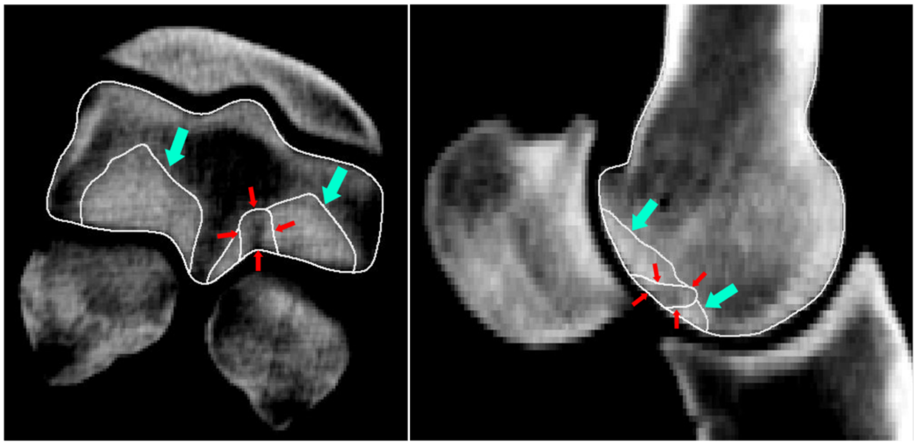

**Figure S1.** Regions isolated using local thresholding. The blue arrows indicate the regions of bone with HU ≥ 1,200, which was qualitatively determined to encompass sclerotic bone. The red arrows indicate the region of bone with HU ≤ 1,200, which was qualitatively determined to encompass damaged bone associated with SBI.

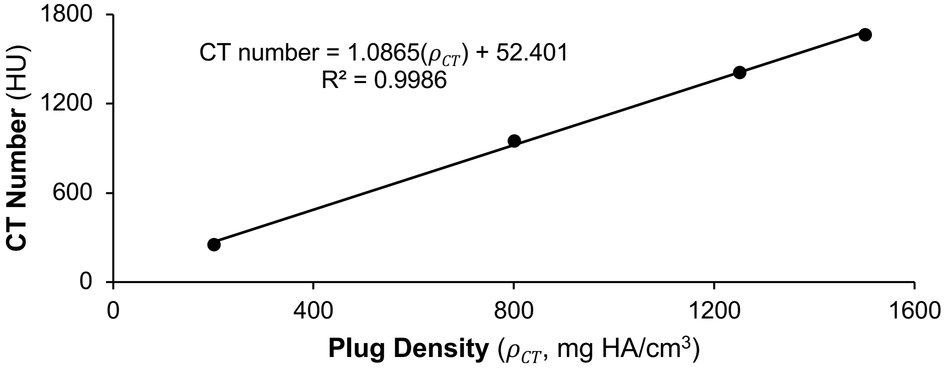


**Figure S2.** Calibration of the CT number (HU) to phantom plug density.

$$\rho_{CT} = (CT\ number - 52.4010)/1.0865 \qquad (equation\ S1)$$

**Figures S3 to S9** show sCT images and the FE-predicted strain plots for Horses #1-3, #5, #6, #7. Frontal oblique (**a**), sagittal (**b**), and transverse oblique (**c**) views of the sCT images are shown. The 3D model of the distal MC3 bone was created with the regions of high sclerosis and lysis (if present) separated (**d**). First principal strain is shown in the frontal oblique (**e**), sagittal (**f**), and transverse oblique (**g**) slices, as well as the palmar joint surface (**h**). Flexed dorsopalmar digital radiograph of the fetlock, if available (**i**) LF – left thoracic limb, RF – right thoracic limb. **Note**: Image montages from Horse #2 LF and #4 LF are presented in Figure 2 in the main manuscript.

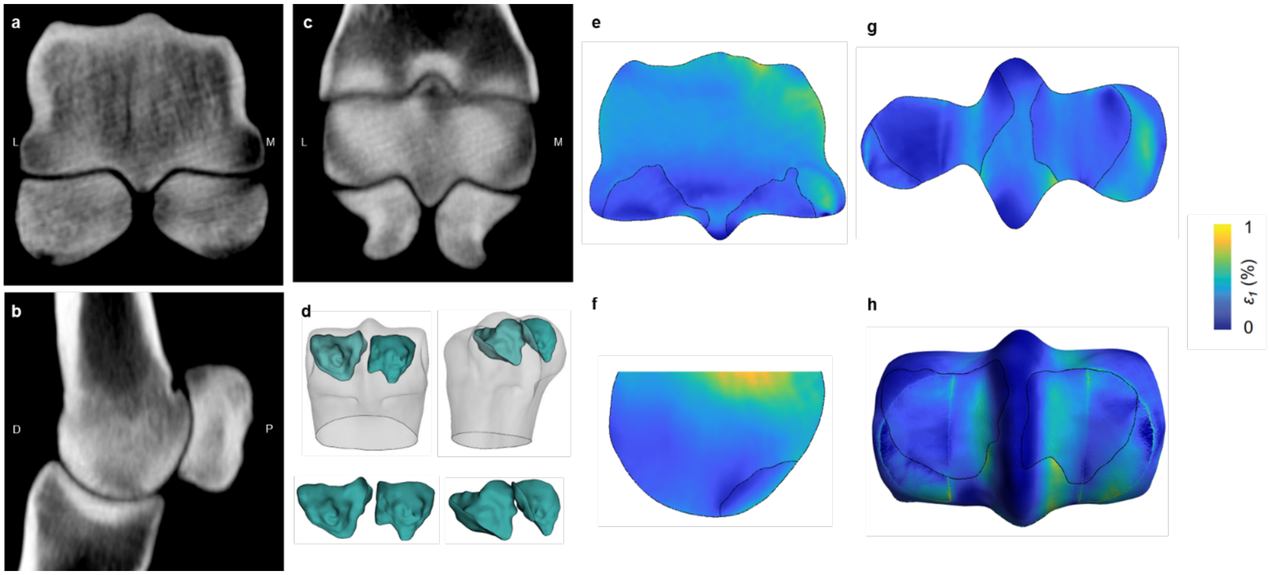


**Figure S3**. Horse #1 LF.

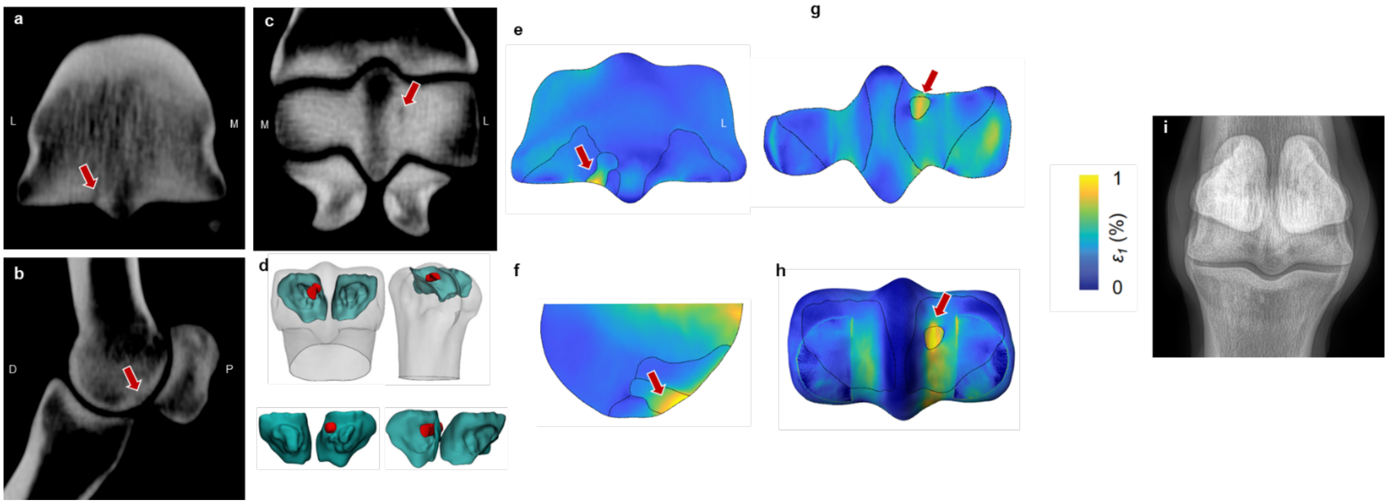


**Figure S4**. Horse #2 RF.

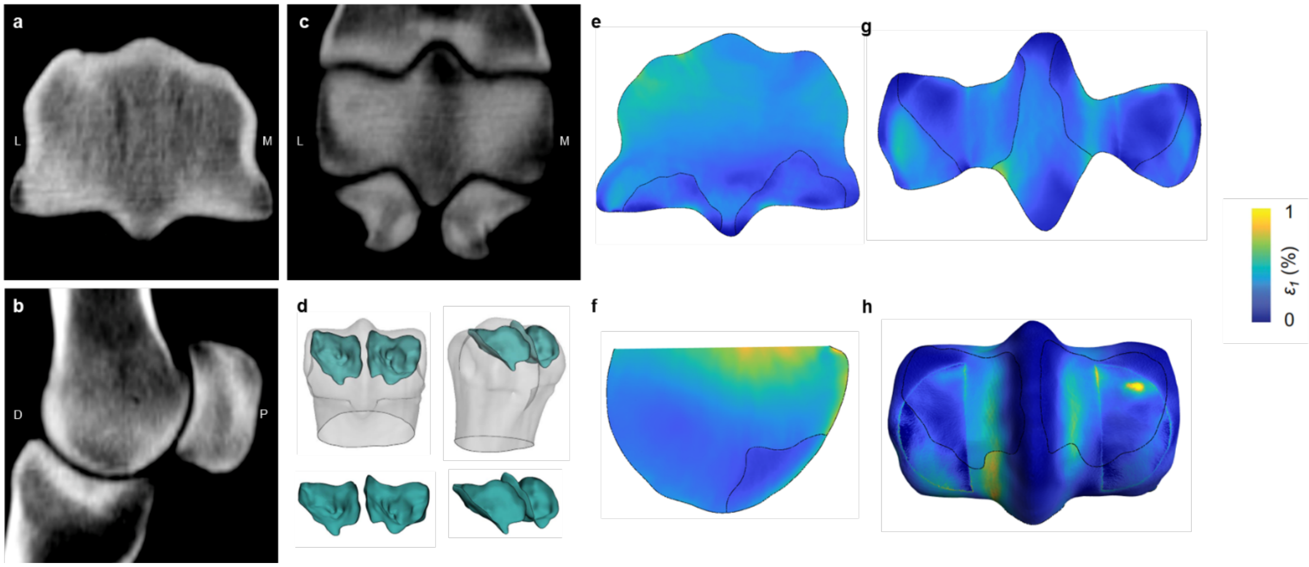


**Figure S5**. Horse #3 RF.

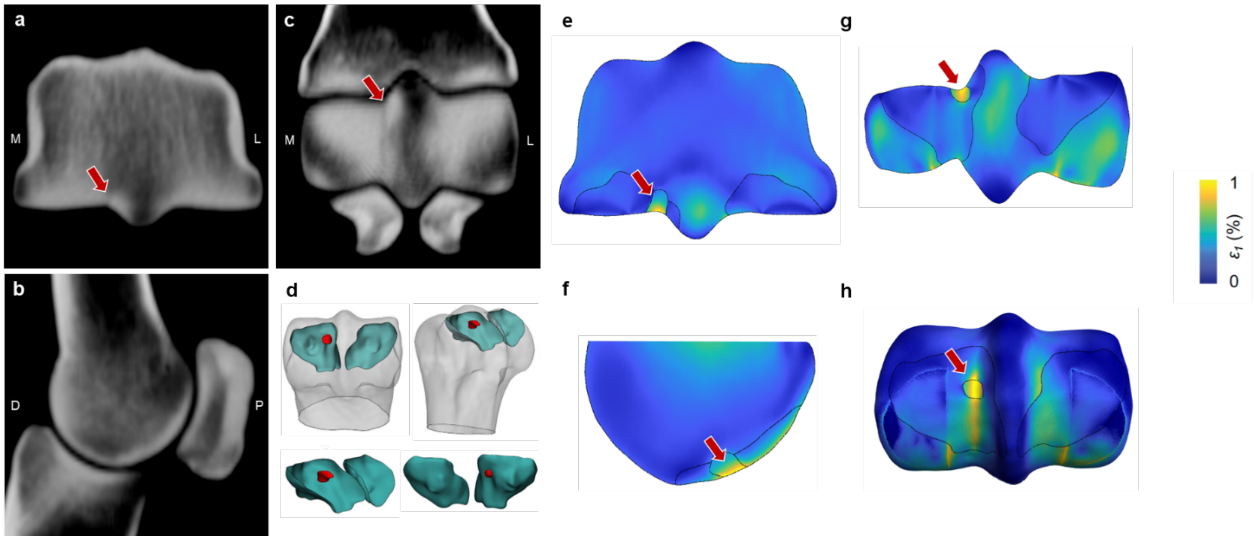


**Figure S6**. Horse #5 LF.

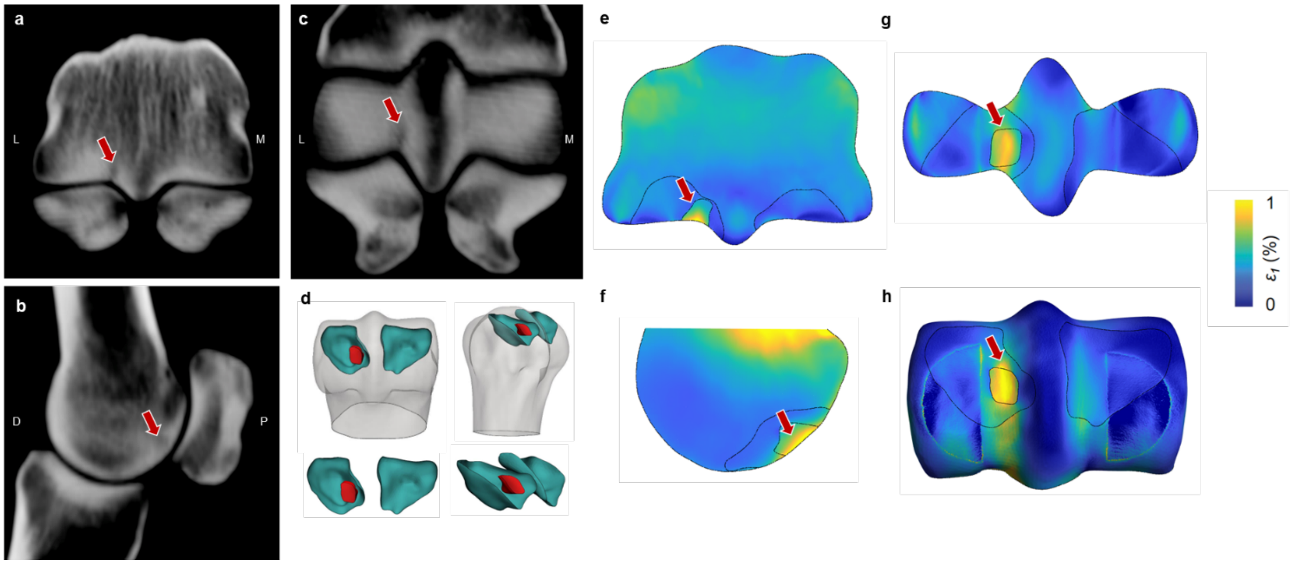


**Figure S7**. Horse #5 RF.

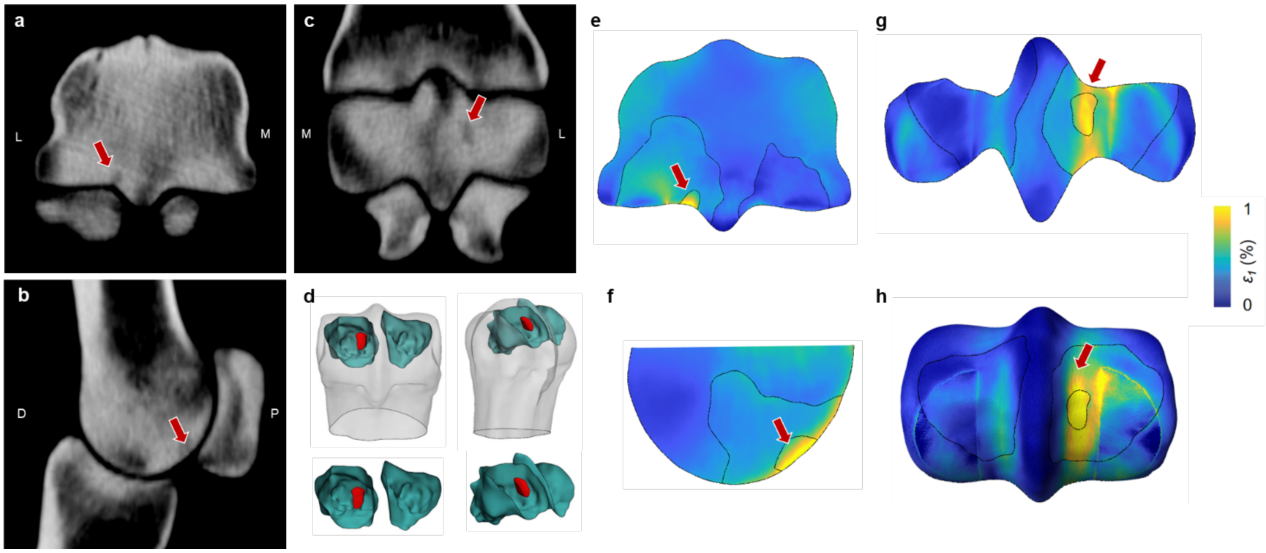


**Figure S8**. Horse #6 RF.

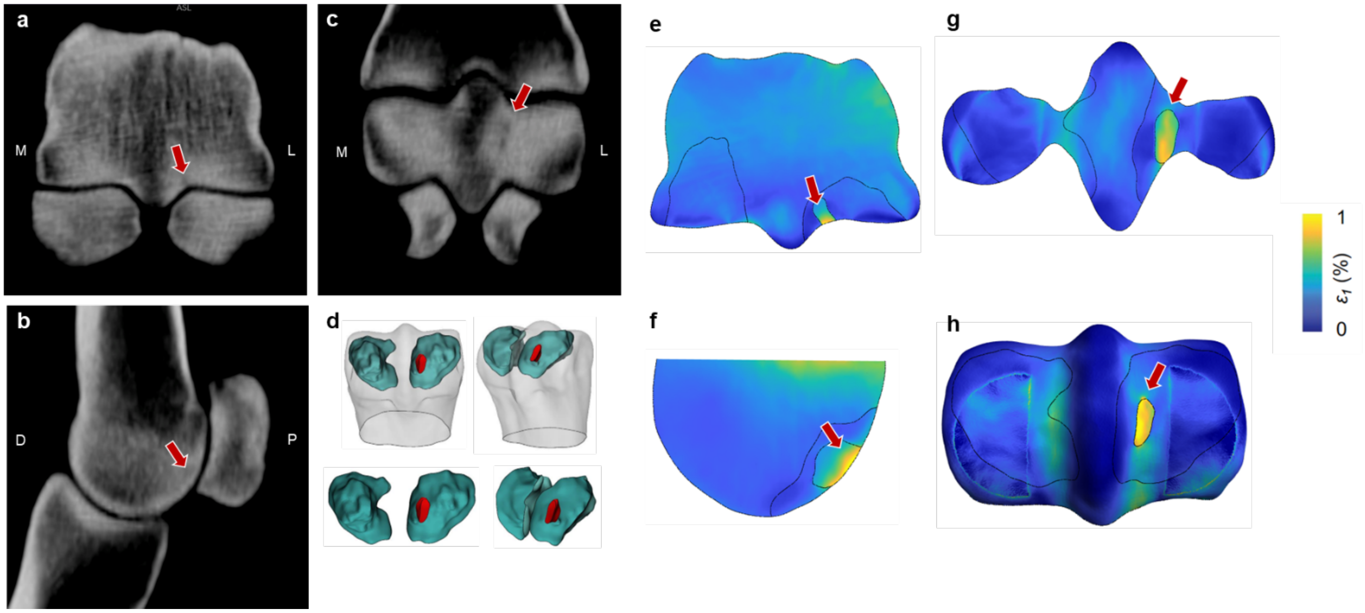

**Figure S9**. Horse #7 LF.

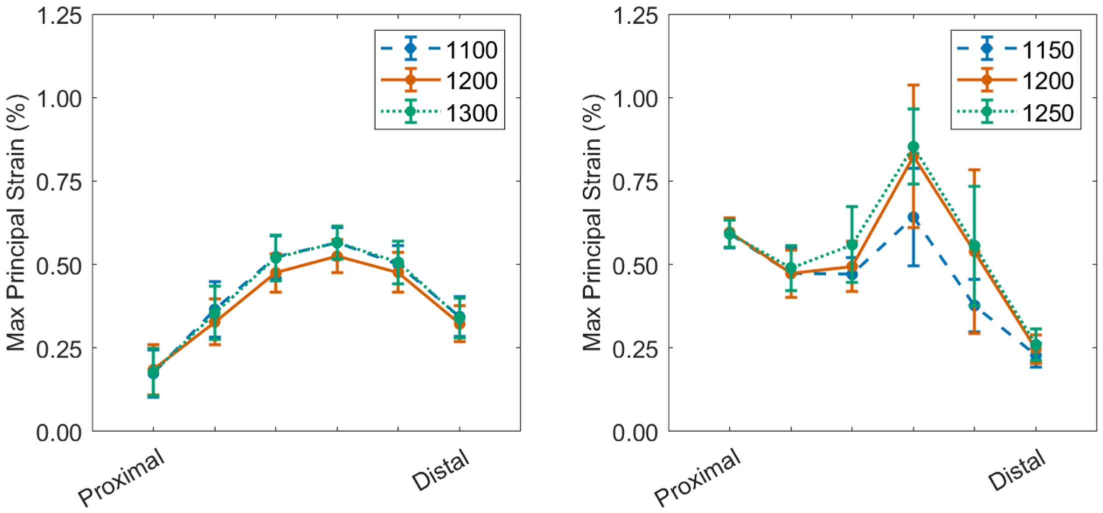


**Figure S10.** Sensitivity of FE-predicted mean PSG first principal strain in the medial (left, CTRL) and lateral (right, CASE) condyles of Horse #7 LF to the three different thresholds used for segmentation of the sclerotic (left, CTRL) and lytic regions.

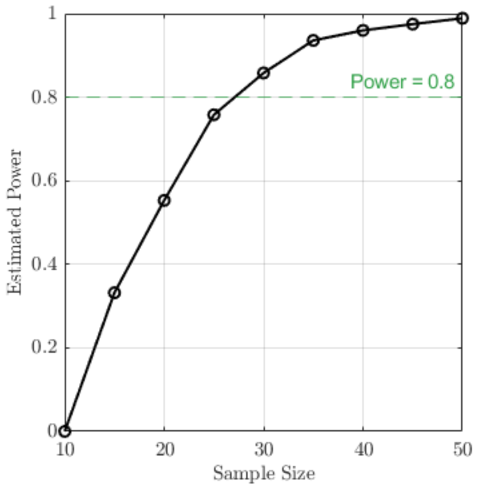

**Figure S11.** The classifier's power is expected to reach 0.8 at a sample size of n=30.
